# Variation in linked selection and recombination drive genomic divergence during allopatric speciation of European and American aspens

**DOI:** 10.1101/029561

**Authors:** Jing Wang, Nathaniel R. Street, Douglas G. Scofield, Pär K. Ingvarsson

## Abstract

Despite the global economic and ecological importance of forest trees, the genomic basis of differential adaptation and speciation in tree species is still poorly understood. *Populus tremula* and *P. tremuloides* are two of the most widespread tree species in the Northern Hemisphere. Using whole-genome re-sequencing data of 24 *P. tremula* and 22 *P. tremuloides* individuals, we find that the two species diverged ∼2.2-3.1 million years ago, coinciding with the severing of the Bering land bridge and the onset of dramatic climatic oscillations during the Pleistocene. Both species have experienced substantial population expansions following long-term declines after species divergence. We detect widespread and heterogeneous genomic differentiation between species, and in accordance with the expectation of allopatric speciation, coalescent simulations suggest that neutral evolutionary processes can account for most of the observed patterns of genomic differentiation. However, there is an excess of regions exhibiting extreme differentiation relative to those expected under demographic simulations, which is indicative of the action of natural selection. Overall genetic differentiation is negatively associated with recombination rate in both species, providing strong support for a role of linked selection in generating the heterogeneous genomic landscape of differentiation between species. Finally, we identify a number of candidate regions and genes that may have been subject to positive and/or balancing selection during the speciation process.

## Introduction

Understanding how genomes diverge during the process of speciation has received a great deal of attention in the evolutionary genetics literature in recent years (Nosil et al. 2009; Strasburg et al. 2012; Seehausen et al. 2014). Under strict neutrality, differentiation is expected to accumulate as a result of the stochastic fixation of polymorphisms by genetic drift (Coyne and Orr 2004). Demographic processes, such as population bottlenecks or expansions, can accelerate or decelerate the rate of differentiation through changes in the effective population sizes of nascent daughter species (Avise 2000). Random genetic drift and demographic processes are both expected to affect the entire genome (Luikart et al. 2003). Natural selection, however, only influence loci involved in ecological specialization and/or reproductive isolation, resulting in patterns of polymorphisms and divergence that deviate from neutral predictions (Luikart et al. 2003; Via 2009). The functional architectures of genomes, e.g. mutation and recombination rates, are also important factors in determining genomic landscape of differentiation (Noor and Bennett 2009; Nachman and Payseur 2012; Renaut et al. 2013). For example, suppressed recombination could increase genetic differentiation either by limiting inter-species gene flow to prevent the breakup of co-adapted alleles, or through the diversity-reducing effects of linked selection (Noor and Bennett 2009). However, disentangling the relative importance of these evolutionary forces when interpreting patterns of genomic divergence remains a challenge in speciation genetics.

With the advance of next generation sequencing (NGS) technologies, a growing number of studies have found highly heterogeneous patterns of genomic differentiation between recently diverged species (Turner et al. 2005; Ellegren et al. 2012; Renaut et al. 2013; Carneiro et al. 2014; Feulner et al. 2015). A common explanation for these patterns is that levels of gene flow between species differ across the genome. Increased genetic divergence is usually observed in a small number of regions containing loci involved in reproductive isolation (‘speciation islands’), where as the remainder of the genome is still permeable to ongoing gene flow and therefore shows lower levels of differentiation (Nosil et al. 2009; Sousa and Hey 2013). However, some recent studies have argued that highly differentiated regions represent ‘incidental islands’ that are not tied to the speciation processes per se. Rather they are seen simply as a result of the diversity-reducing effects of linked selection that accelerate lineage sorting of ancestral variation and increase interspecific differentiation, especially in regions of reduced recombination (Turner and Hahn 2010; Cruickshank and Hahn 2014). In addition, long-term balancing selection is supposed to maintain stable trans-species polymorphisms and leaves signatures of unusually low genetic differentiation between species (Charlesworth 2006). Under these scenarios, natural selection alone is sufficient to generate patterns of heterogeneous genomic differentiation even under complete allopatry (Noor and Bennett 2009; Turner and Hahn 2010). Finally, strictly neutral forces, such as stochastic genetic drift and complex demographic processes, can also create heterogeneous genomic divergence and generate patterns of divergence and polymorphism that mimic the effects of selection (Nosil et al. 2009; Campagna et al. 2015). In general, the three hypotheses listed above are not mutually exclusive and exhaustive examination of these hypotheses requires detailed information on the speciation process, such as the timing of speciation, the geographic and demographic context in which it occurred (Nosil and Feder 2012).

Although largely understudied compared to other model species, forest trees represent a promising system to understand the genomic basis of species divergence and adaptive evolution; as a group they have developed diverse strategies to adapt and thrive across a wide range of climates and environments (Neale and Kremer 2011). *Populus tremula* (European aspen) and *P. tremuloides* (American aspen) are two of the most ecologically important and geographically widespread tree species of the Northern Hemisphere (Figure 1a). Both are keystone species, display rapid growth, with high tolerance to environmental stresses and long-distance pollen and seed dispersal via wind (Eckenwalder 1996; Müller et al. 2012). In addition, they both harbor among the highest level of intraspecific genetic diversity reported in plant species so far (Wang et al. forthcoming). Based on their morphological similarity and close phylogenetic relationships, they are considered to be sister species, or less commonly, conspecific subspecies (Eckenwalder 1996; Wang et al. 2013). They can readily cross and artificial hybrids usually show high heterosis (Hamzeh and Dayanandan 2004; Tullus et al. 2012). A recent study based on a handful of nuclear and chloroplast loci suggests that the first opening of the Bering land bridge may have driven the allopatric speciation of the two species (Du et al. 2015).

**Figure 1.**
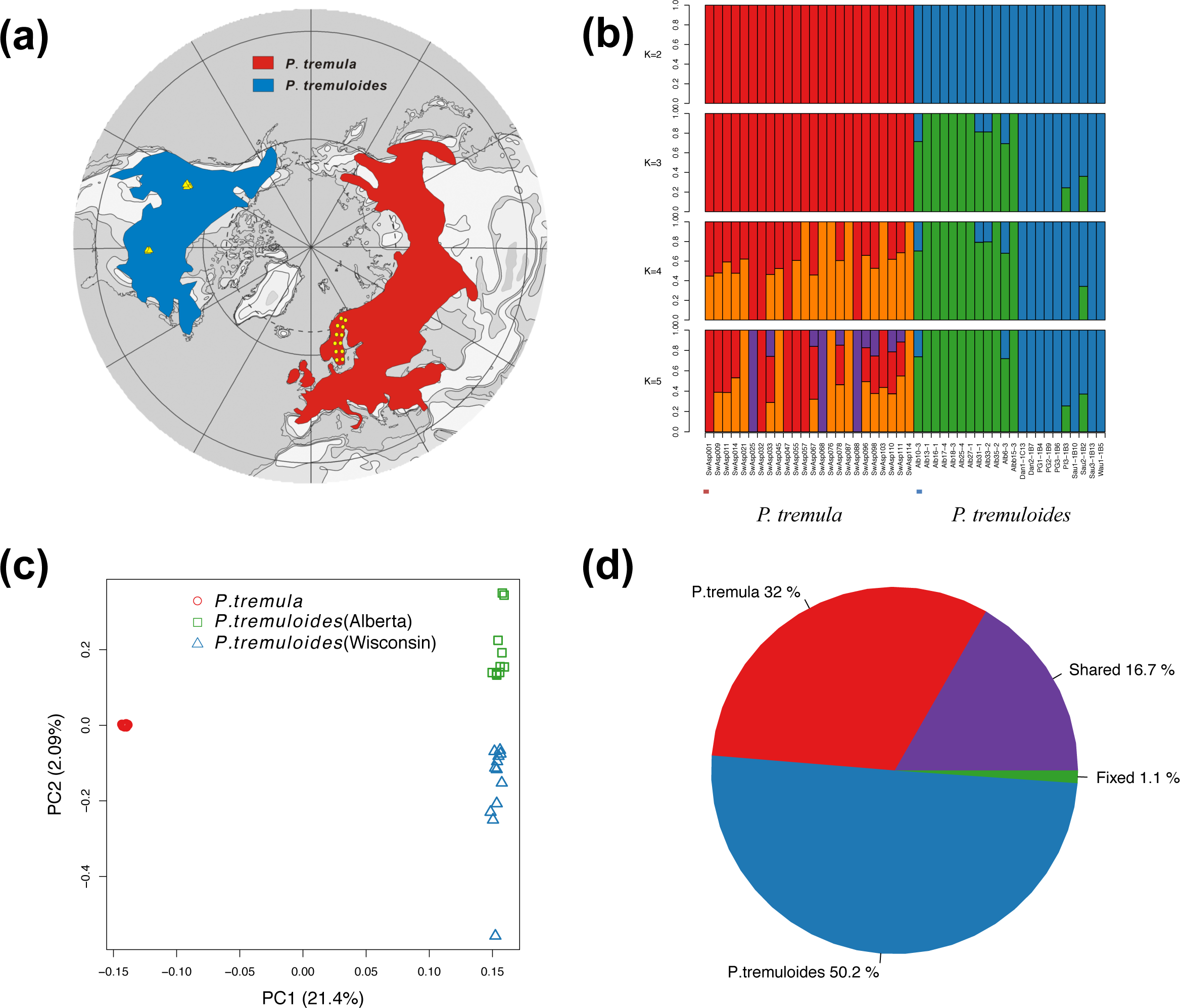
Geographic distribution and genetic structure of 24 *Populus tremula* and 22 *P. tremuloides.* (a) Map showing the current geographic distribution of *P. tremula* (red) and *P. tremuloides* (blue). Yellow circles and triangles indicate the locations where the 24 individuals of *P. tremula* and 22 individuals of *P. tremuloides* were sampled. (b) Genetic structure of the two species inferred using NGSadmix. The y-axis quantifies subgroup membership, and the x-axis shows the sample ID for each individual. (c) Principal component analysis (PCA) plot based on genetic covariance among all individuals of *P. tremula* (red circle) and *P. tremuloides* (green square and blue triangle). The first two principle components (PCs) are shown, with PC1 explaining 21.04% (*P*=2.51×10^−19^, Tacey-Widom test) of the overall genetic variation and separating the two species and PC2 explaining 2.09% (*P*=9.65×10^−4^, Tracy-Widom test) of the overall variation and separating samples from Wisconsin (blue triangle) and Alberta (green square) in *P. tremuloides*. (d) Pie chart summarizing the proportion of fixed, shared and exclusive polymorphisms of the two species.

Due to their continent-wide distributions, extraordinary levels of genetic and phenotypic diversity, along with the availability of a high-quality reference genome in the congener, *P. trichocarpa* (Tuskan et al. 2006), *P. tremula* and *P. tremuloides* represent a promising system for evaluating how various evolutionary processes have shaped the patterns of genomic divergence during speciation. In this study, we use whole-genome re-sequencing data from both species to estimate and infer their divergence time and historical demographic processes of the two species. Explicit characterizations of the demographic history not only allow us to estimate historical population size fluctuations in both species, but also increase the accuracy of identifying regions or genes that have been under natural selection. By incorporating the inferred demographic scenarios into the null model, we investigate the extent to which demographic and selective events have contributed to the overall patterns of genomic differentiation between the two species. We also identify a number of outlier regions and genes that likely have evolved in response to positive and/or balancing selection during the speciation process.

## Results

We generated whole-genome resequencing data for 24 *P. tremula* and 22 *P. tremuloides*. The high extent of conserved synteny between the genomes of aspen and *P. trichocarpa* (Pakull et al. 2009; Robinson et al. 2014) allowed us to map all reads to the *P. trichocarpa* reference genome (v3.0) (Tuskan et al. 2006) after adapter removal and quality trimming (see Materials and Methods). More than 88% of sequenced reads were aligned and the mean coverage of uniquely mapped reads per site was 25.1 and 22.5 in samples of *P. tremula* and *P. tremuloides*, respectively (Table S1). Two complementary bioinformatics approaches were used in this study (Figure S1): (1) For those population genetic statistics that relied on inferred site-frequency-spectrum (SFS), estimation was performed directly from genotype likelihoods without calling genotypes (Nielsen et al. 2011) as implemented in ANGSD (Korneliussen et al. 2014). (2) For those estimations that required accurate genotype calls, single nucleotide polymorphisms (SNPs) and genotypes were called with HaplotypeCaller in GATK (Danecek et al. 2011). In total, we identified 5,894,205 and 6,281,924 SNPs passing filtering criteria (see Materials and Methods) across the 24 *P. tremula* samples and 22 *P. tremuloides* samples, respectively.

## Population structure

We used NGSadmix (Skotte et al. 2013) to infer individual ancestry based on genotype likelihoods, which takes the uncertainty of genotype calling into account. It clearly sub-divided all sampled individuals into two species-specific groups when the number of clusters (*K*) was 2 (Figure 1b). When *K* = 3, there was evidence for further population sub-structuring in *P. tremuloides*, where individuals from populations of Alberta and Wisconsin clustered into two subgroups. With *K* = 4, most individuals of *P. tremula* were inferred to be a mixture of two genetic components, showing slight clinal variation with latitude. No further structure was found when *K* = 5 (Figure 1b). A principal component analysis (PCA) further supported these results (Figure 1c). Only the first two components were significant based on the Tracy-Widom test (Table S2), which explained 21.4% and 2.1% of total genetic variance, respectively (Figure 1c). Among the total number of polymorphisms in the two species, fixed differences between *P. tremula* and *P. tremuloides* accounted for 1.1%, whereas 16.7% of polymorphisms were shared between species, with the remaining polymorphic sites being private in either of the two species (Figure 1d).

To further examine the extent of population subdivision in *P. tremuloides*, we measured *F*_ST_ and d_xy_ between the two subpopulations (Alberta and Wisconsin) along individual chromosomes (Table S3). We found low levels of genetic differentiation (average *F*_ST_: 0.0443±0.0325) between the two subpopulations (Table S3). Total sequence differentiation in the inter-population comparison (mean dxy = 0.0165±0.0083) was similar to mean sequence differences in intra-population comparisons (π_Alberta_: 0.0161±0.0081; π_Wisconsin_: 0.0157±0.0080, Table S3), indicating that individuals of the two populations were genetically not more different from each other than individuals within each population. Based on the summaries of site frequency spectrum (Tajima’s *D* and Fay & Wu’s *H*), both populations exhibited strong skews toward low-frequency variants (negative *D*) and intermediate skews toward high frequency-derived variants (negative *H*) (Table S3), suggesting that they likely experienced similar species-wide demographic events.

## Demographic histories

We used *fastsimcoal2* (Excoffier et al. 2013), a coalescent simulation-based method, to infer the past demographic histories of *P. tremula* and *P. tremuloides* from the joint site frequency spectrum. Eighteen divergence models were evaluated (Figure S2; Table S4), and all models began with the split of the ancestral population into two derived populations and differed in terms of (i) whether post-divergence gene flow was present or not, (ii) levels and patterns of gene flow between the two species, and (iii) how population size changes occurred, either at the time of species divergence or afterwards (Figure S2). The best-fitting model was a simple isolation-with-migration model where, after the two species diverged, *P. tremuloides* experienced exponential growth and whereas a stepwise population size change occurred in *P. tremula* (Figure 2a). The exact parameter estimates of divergence time, gene flow, population sizes and their associated 95% confidence intervals (CIs) are given in Table 1. The estimated divergence time between *P. tremula* and *P. tremuloides* (T_DIV_) was ∼2.3 million years ago (Mya) (bootstrap range [BR]: 2.2-3.1 Mya). The contemporary effective population sizes (*N*_e_) of *P. tremula* (N_*P.tremula*_) and *P. tremuloides* (N_*P.tremuloides*_) were 102,814 (BR: 93,688-105,671) and 309,500 (BR: 247,321-310,105) respectively, with both being larger than the effective population size of their common ancestor (N_ANC_ = 56,235 [48,012-69,492])). Gene flow (2*N*_e_m, where *N*_e_ is the effective population size and m is the migration rate) from *P. tremuloides* to *P. tremula* was higher (0.202 migrants per generation [0.156-0.375]) than in the opposite direction (0.053 [0.052-0.117]), most likely reflecting the higher *N*_e_ in *P. tremuloides* than in *P. tremula* (Slatkin 1985). Overall, the migration rates in both directions were fairly low given the large *N*_e_ of both species (Morjan and Rieseberg 2004), which is not unexpected given the large geographical distance and disjunct distributions between the two species.

**Table 1.**
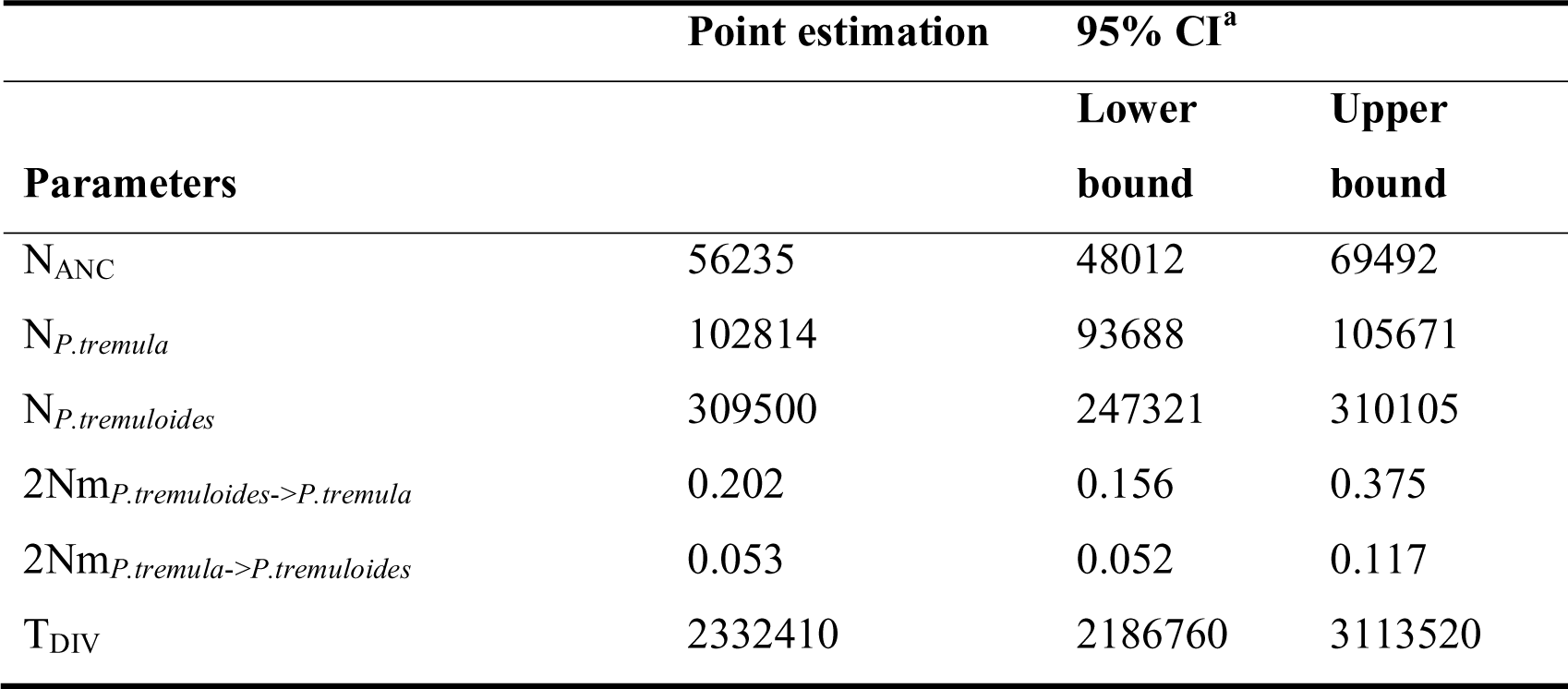
Inferred demographic parameters of the best-fitting demographic model shown in Figure 2a. Parameters are defined in Figure 2a. N indicates the effective population size of *P. tremula*, *P. tremuloides* or their ancestral population, m indicates the migration rates between species on either direction, TDIV indicates the estimated divergence time between the two species obtained from *fastsimcoal2*. ^a^Parametric bootstrap estimates obtained by parameter estimation from 100 data sets simulated according to the overall maximum composite likelihood estimates shown in point estimation columns. Estimations were obtained from 100,000 simulations per likelihood.

**Figure 2.**
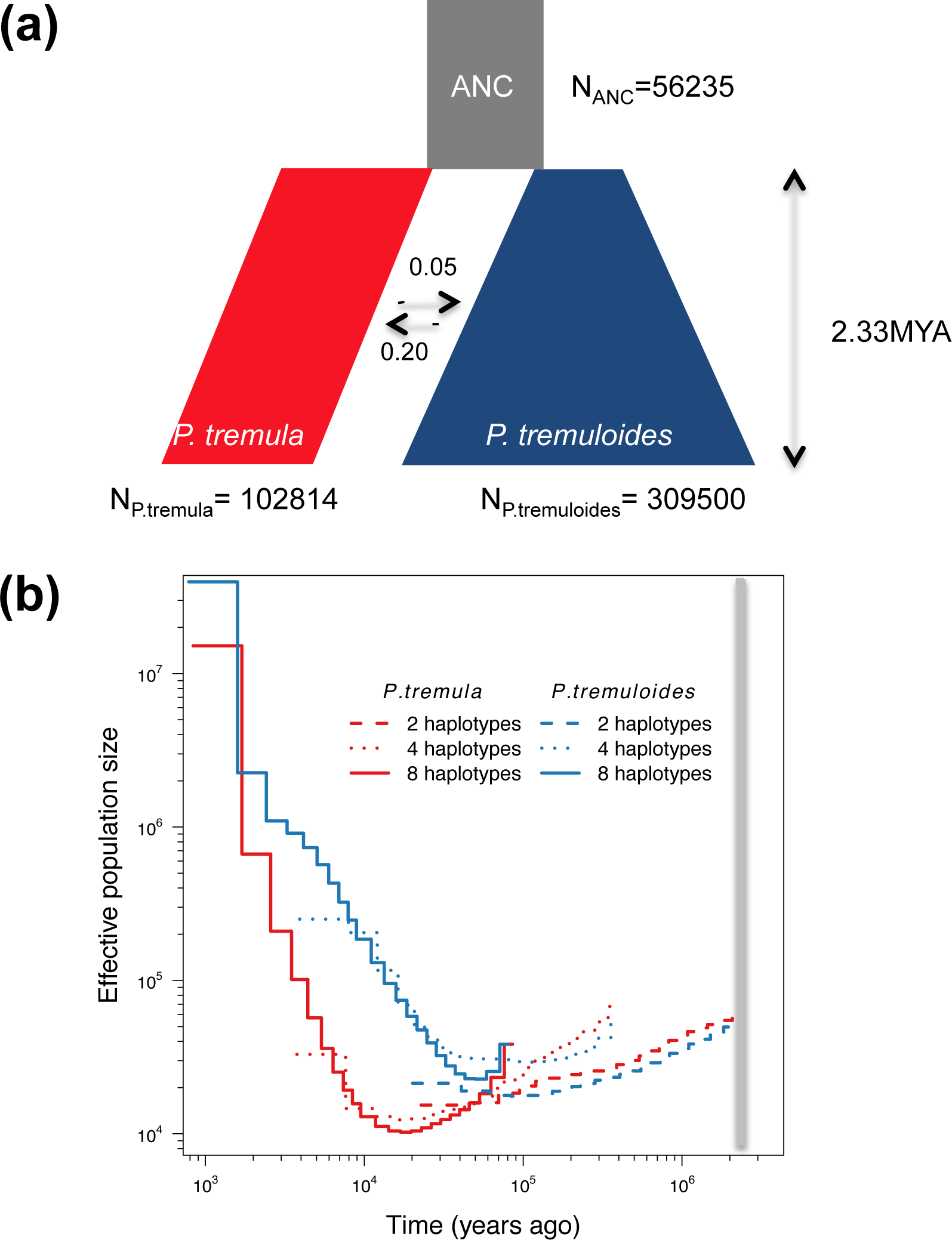
Demographic history of *Populus tremula* and *P. tremuloides.* (a) Simplified graphical summary of the best-fitting demographic model inferred by *fastsimcoa12*. (b) Multiple sequential Markovian coalescent (MSMC) estimates of the effective population size (N_e_) changes for *P. tremula* (red line) and *P. tremuloides* (blue line) based on the inference from two (dashed), four (dotted) and eight (solid) phased haplotypes in each species. Time scale on the x-axis is calculated assuming a neutral mutation rate per generation (μ) = 3.75×10^−8^ and generation time (g) = 15 years. The grey bar indicates the speciation time inferred by *fastsimcoal2*.

We employed the multiple sequential Markovian coalescent (MSMC) method to investigate changes of *N*_e_ over time based on inferring the time to the first coalescence between pairs of haplotypes (Schiffels and Durbin 2014). Higher resolution of recent population size changes is expected when more haplotypes are used (Schiffels and Durbin 2014). We therefore applied MSMC to phased whole-genome sequences from one (two haplotypes), two (four haplotypes) and four (eight haplotypes) individuals in each species, respectively. We did not include more haplotypes because of the high computational cost of larger samples. The MSMC-based estimates of *N*_e_ for both *P. tremula* (60,796) and *P. tremuloides* (49,701) at the beginning of species divergence (around 2.3 Mya) were very similar to the *fastsimcoa12*-based estimates of *N*_e_ for their ancestral population (Figure 2). The two species experienced similar magnitudes of population decline following their initial divergence (Figure 2b). Population expansion in *P. tremuloides* occurred around 50,000-70,000 years ago and continued up to the present (Figure 2b), whereas *P. tremula* experienced a population expansion following a substantially longer periods of bottleneck (Figure 2b).

To assess the possible confounding effects of population subdivision and biased sampling scheme on demographic inferences in both species, we first applied MSMC analysis using *P. tremuloides* individuals originating from populations in Alberta and Wisconsin separately and compared them to the result obtained from the pooled samples (Figure S3). Although the Wisconsin population was found to have undergone a decline in population size during the last 2000-3000 years ago (Figure S3), both local populations of *P. tremuloides* show evidence for a longer period of species-wide expansion when compared with *P. tremula*, which was in accordance with the results observed from the pooled samples. Second, demographic inferences of both species were also supported by the summary statistics based on the nucleotide diversity (θ_π_) and site frequency spectrum (Tajima’s *D* and Fay & Wu’s *H*) (Figure S4). The θ_π_ in the two subpopulations of *P. tremuloides* were all marginally higher than in *P. tremula*, suggesting that the large effective population size found in *P. tremuloides* is not influenced by the presence of intraspecific population subdivision (Figure S4a). In addition, the signal of more negative values of Tajima’s *D* in both local and pooled samples of *P. tremuloides* (Figure S4b) suggest that it may have gone through a more pronounced and/or longer period of population expansion compared to *P. tremula*. The lower values of the genome-wide Fay & Wu’s *H* in *P. tremula* (Figure S4c), on the other hand, might reflect the relatively longer period of low population size during the bottleneck. Taken together, these results suggest that population subdivision of *P. tremuloides* and the unbalanced sampling schemes between the two species have negligible effects on our demographic inferences.

## Genome-wide patterns of differentiation and identification of outlier regions against the best-fitting demographic model

To investigate patterns of interspecific genetic differentiation across the genome, we calculated the standard measure of genetic divergence, *F*_ST_, between *P. tremula* and *P. tremuloides* over non-overlapping 10 Kbp windows (Figure 3). Levels of genetic differentiation varied greatly throughout the genome, with the majority of windows showing high genetic differentiation (mean *F*_ST_ = 0.386) between species (Figure 3).

**Figure 3.**
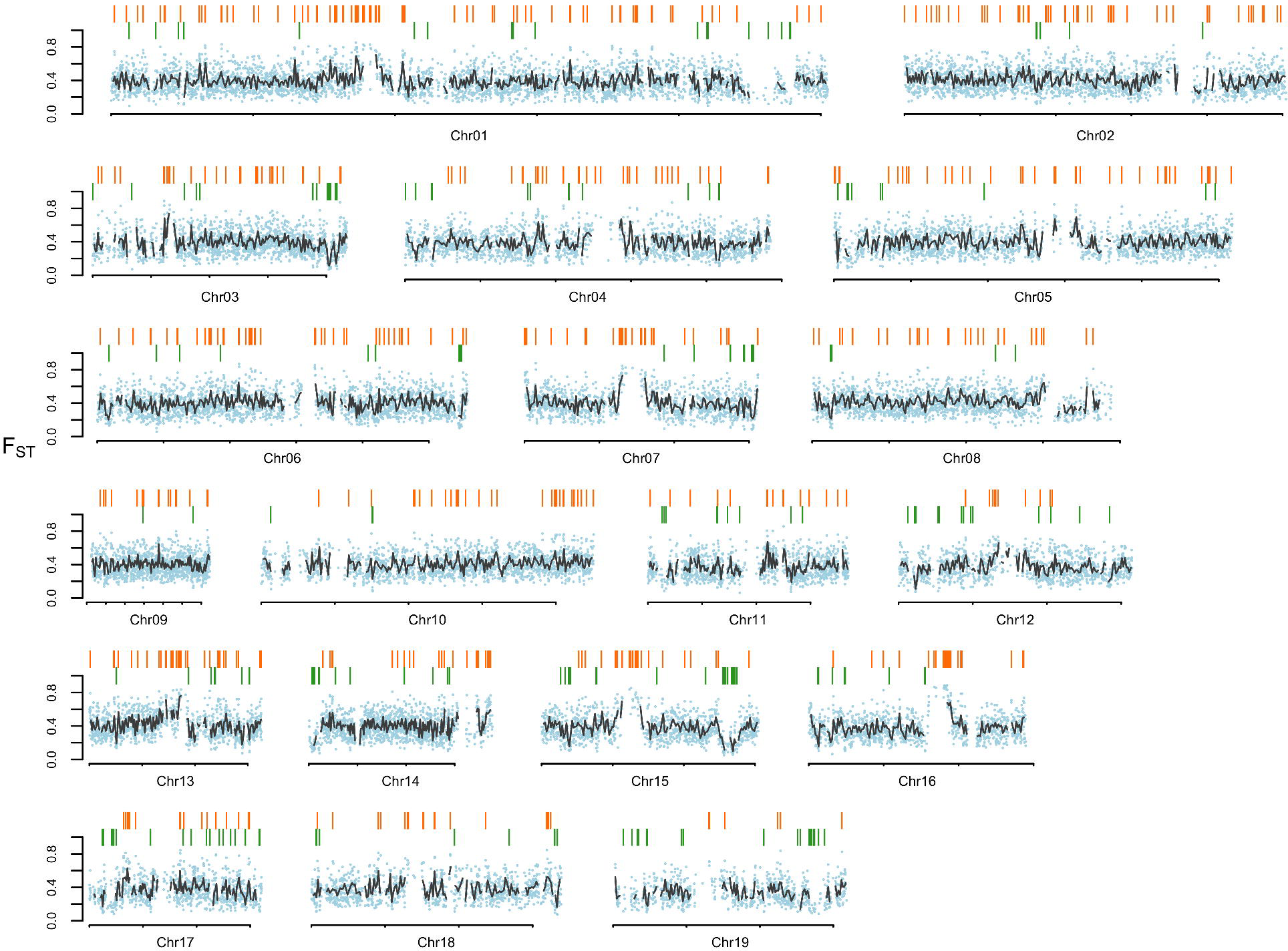
Genome-wide divergence. Chromosomal distribution of genetic differentiation (*F*_ST_) between *Populus tremula* and *P. tremuloides*. The small, light blue dots indicate *F*_ST_ values estimated over 10 Kbp non-overlapping windows. Grey lines indicate *F*_ST_ values estimated over 100 Kbp non-overlapping windows. Locations for windows displaying extreme differentiation relative to demographic simulations are highlighted with colored bars above the plot. Among them, candidate windows displaying significantly high differentiation (orange bars) are located on the topside; candidate windows displaying significantly low differentiation (green bars) are located at the bottom.

In order to test the extent to which historical demographic events can explain the observed patterns of genetic divergence between the two species, we used coalescent simulations performed in *msms* (Ewing and Hermisson 2010) to compare the observed patterns of differentiation to that expected under three demographic models (Figure S5). The demographic scenario in model 1 was same as the best-fitting model inferred by *fastsimcoa12* (Figure S5a; Table S5). In another two models, we incorporated the population subdivision of *P. tremuloides* into the best-fitting demographic model. In model 2 (Figure S5b; Table S5), we assume there was no gene flow between the two subpopulations of *P. tremuloides* and explored different values of their divergence time until the simulated *F*_ST_ values between the two subpopulations matched those observed (Figure S6). The same procedure was applied to model 3 (Figure S5c; Table S5), except that we there assume the per-generation gene flow between the two subpopulations of *P. tremuloides* (4*N*_e_m) was equal to 10 and increased their divergence time in tandem with gene flow. To assess the fit of these models, we compared two summary statistics, θ_π_ and Tajima’s D, between the simulated and observed data for both species. As can be seen from Figure S7 and Figure S8, there was generally a good agreement between observed and simulated data sets for all three models. In addition, the above three models showed consistent distributions of simulated *F*_ST_ values between *P. tremula* and *P. tremuloides*, indicating that the presence of population subdivision in *P. tremuloides* has little effect on the overall patterns of genomic divergence that we observe between the two species (Figure S9).

Comparing the observed distribution of inter-specific *F*_ST_ with that obtained from simulations based on the best-fitting demographic model, we found that the observed distribution was flatter and contained greater proportions of regions falling in the extremes of distribution (Figure 4a). We used the distribution of inter-specific *F*_ST_ based on 500,000 coalescent simulations to identify outlier windows that were likely targets of natural selection by performing a false discovery rate (FDR) of 1% as our cut-off. With this approach, 674 and 262 windows (FDR<0.01) were, respectively, identified as showing exceptionally high and low *F*_ST_ between the two species (Figure 4a). After examining the genomic distribution, physical sizes and overlaps of these outlier windows, we found that both highly and lowly differentiated regions were randomly distributed across the genome (Figure 3) and that the sizes of these regions appeared to be rather small, with the majority occurring on a physical scale smaller than 10 Kbp (Figure S10).

**Figure 4.**
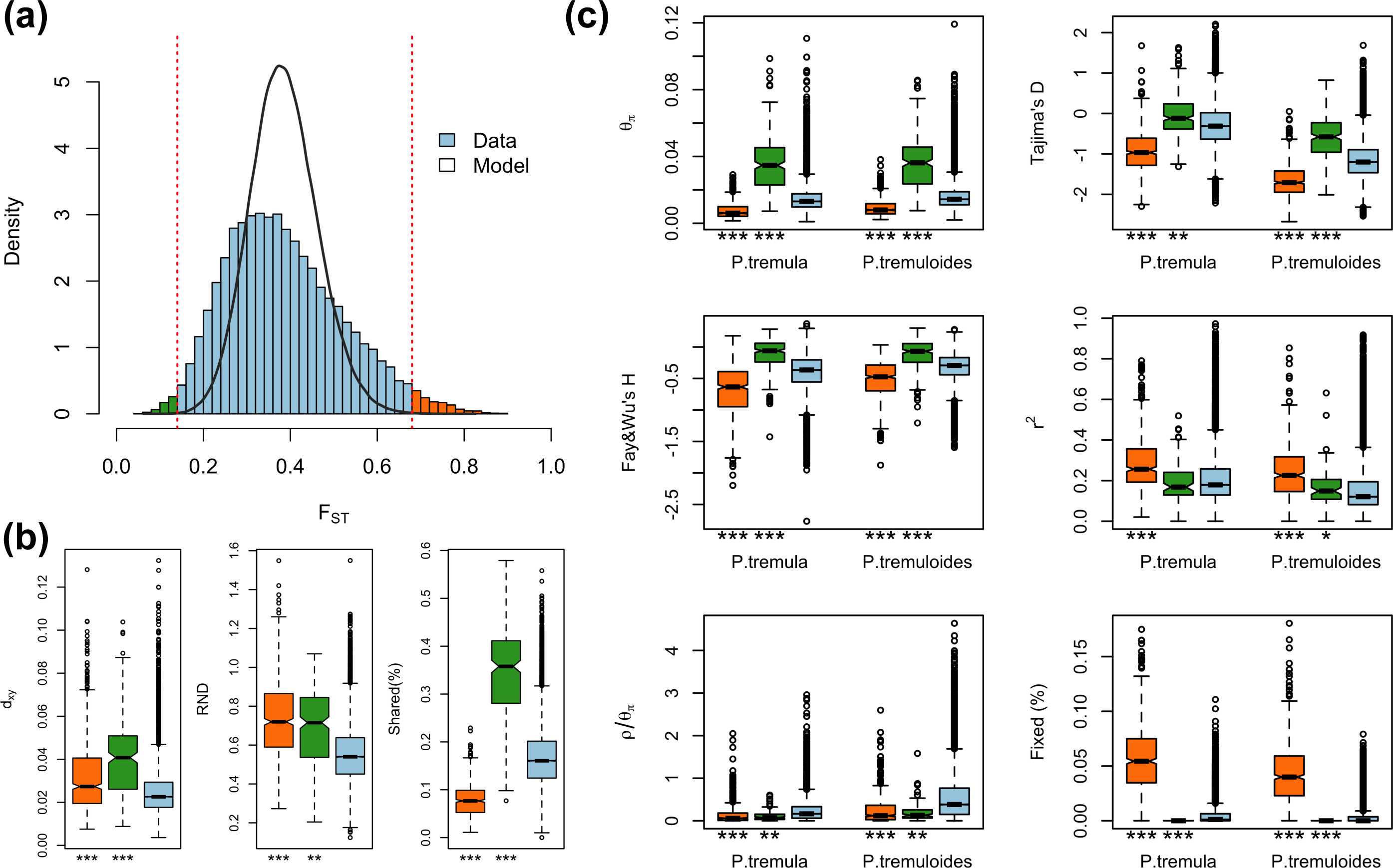
Identification of outlier windows that are candidates for being affected by natural selection. (a) Distribution of genetic differentiation (*F*_ST_) between *P. tremula* and *P. tremuloides* from the observed (blue bar) and simulated data sets (black line). The dashed lines indicate the thresholds for determining significantly (False Discovery Rate <1%) high (orange bars) and low (green bars) inter-specific differentiation based on coalescent simulations. (b) Comparisons of d_xy_, relative node depth (RND) and the proportion of inter-specific shared polymorphisms among regions displaying significantly high (orange boxes) and low (green boxes) differentiation versus the genomic background (blue boxes). (c) Comparisons of multiple population genetic statistics, nucleotide diversity (θ_π_), Tajima’s D, Fay &Wu’s H, linkage disequilibrium (*r*^2^), recombination rate 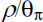, the proportion of fixed differences caused by derived alleles fixed in either *P. tremula* or *P. tremuloides*, among regions displaying significantly high (orange boxes) and low differentiation (green boxes) versus the genomic background (blue boxes). Asterisks designate significant differences between the outlier and the rest of genomic regions by MannWhitney U test (* *P*-value < 0.05; ** *P*-value < le-4; \*\*\**P*-value <2.2e-16).

## Signatures of selection in outlier regions

As *F*_ST_ is a relative measure of differentiation and is thus sensitive to any processes that alter intra-species genetic variation (Charlesworth 1998; Cruickshank and Hahn 2014), we quantified and compared inter-specific genetic differentiation between two unions of outlier windows and the rest of the genome using three additional approaches: (i) pairwise nucleotide divergence between species (d_xy_), which is a measure that is independent of within-species diversity (Nei 1987); (ii) relative node depth (RND) (Feder et al. 2005), which takes into account possible variation in the mutation rate among genomic regions by dividing d_xy_ of the two aspen species with d_xy_ between aspens and a third more distantly related species (*P. trichocarpa*); and (iii) the proportion of inter-specific shared polymorphisms. Compared to the genomic background average, both d_xy_ and RND revealed much greater divergence between the two species in regions of high differentiation (Figure 4b; Table S6) and, in accordance with these patterns, the proportion of inter-specific shared polymorphisms was significantly lower in these regions (Figure 4b; Table S6). In addition, these regions are characterized by multiple signatures of positive selection within one or both species, including significantly reduced levels of polymorphism (θ_π_), skewed allele frequency spectrum towards rare alleles (more negative Tajima’s D), increased high-frequency derived alleles (more negative Fay & Wu’s H), and stronger signals of linkage disequilibrium (LD) (P<0.001, Mann-Whitney U test) (Figure 4c; Table S6). Relative to genome-wide averages, these regions also contained significantly higher proportions of fixed differences that were caused by derived alleles fixed in either species (Figure 4c; Table S6).

In contrast to patterns found in regions of high differentiation, regions of low differentiation showed significantly higher levels of polymorphism, excesses of intermediate-frequency alleles (higher Tajima’s D and Fay & Wu’s H values), higher proportions of inter-specific shared polymorphisms and negligible proportions of fixed differences compared to the genomic background (Figure 4b, c; Table S6). It is therefore likely that some of these regions have been targets of long-term balancing selection in both species (Charlesworth 2006). Consistent with this prediction, we found slightly lower or comparable levels of LD in these regions (Figure 4c; Table S6), which is likely due to the long-term effects of recombination on old balanced polymorphisms (Leffler et al. 2013). The higher dxy and RND values we observe in these regions may, however, be a consequence of the higher levels of ancestral polymorphisms that were maintained predating the split of the two species (Figure 4b; Table S6) (Cruickshank and Hahn 2014).

## Impact of recombination rate on patterns of genetic differentiation

We examined relationships between the scaled recombination rates 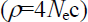 and levels of inter-species divergence over non-overlapping 10 Kbp windows across the genome (Figure S11). We found a significant negative correlation between relative divergence, measured as *F*_ST_ that depends on genetic diversity within species, and recombination rates in both *P. tremula* (Spearman’s ρ=-0.121, *P*-value<0.001) and *P. tremuloides* (Spearman’s ρ=-0.157, P-value<0.001) (Figure S11a). In contrast to *F*_ST_, we observed significantly positive correlations between absolute divergence d_xy_ and recombination rates in both species (*P. tremula:* Spearman’s ρ=0.199, P-value<0.001; *P. tremuloides:* Spearman’s ρ=0.140, *P*-value<0.001) (Figure S11b).

Because 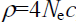, where *c* is the per-generation recombination rate and *N*_e_ is the effective population size, a reduction of *N*_e_ in regions linked to selection will lower local estimates of *ρ* even if local *c* is identical to other regions in the genome. In order to account for such effects and to obtain a measure of recombination that is independent of local *N*_e_, we compared 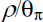 between regions with extreme genetic differentiation and the remainder of the genome. Relative to the genomic background, our results showed significantly suppressed recombination in outlier regions displaying either exceptionally high or low inter-specific differentiation (Figure 4c).

## Genes under selection

The availability of the annotated *P. trichocarpa* genome enabled functional analyses of candidate target genes within regions that were likely under selection. In total, 722 and 391 genes were located in outlier windows displaying exceptionally high and low differentiation (Table S7 and S8), respectively. Compared to the genome overall, we did not find significantly higher gene density in these outlier windows (P>0.05, Mann-Whitney U test; Figure S12). We used the Gene Ontology (GO) assignments of those candidate genes to assess whether specific GO terms were significantly over-represented. After accounting for multiple comparisons, we did not detect over-representation of any functional category among the candidate genes within regions of high differentiation. However, we identified 60 significantly overrepresented GO terms for genes located within regions showing significantly low genetic differentiation and that were likely candidates for being under the influence of balancing selection. Most of these GO categories were associated with immune and defense responses, signal transduction or apoptosis (Table S9). Nevertheless, some caution should be applied when interpreting these results since we observed a skewed pattern of low coverage breadth in outlier windows displaying significantly low differentiation compared to either the genomic background or to those highly differentiated windows (Figure S13). Such unequal coverage breadth likely results from the inherent technical hurdle of short-read sequencing technologies and likely represents difficulties of mapping short reads to a reference genome in highly polymorphic regions (Brandt et al. 2015). After stringent quality filtering, more reads were discarded in these regions, which further decreased the amount of usable information. Therefore, future studies, incorporating a combination of careful experimental design and long-read sequencing technologies, are needed to evaluate the accuracy of the evolutionary significance of balancing selection found in the candidate genes shown here.

## Discussion

We use a population genomic approach to resolve the evolutionary histories of two widespread and closely related forest tree species, and to highlight how genome-wide patterns of differentiation have been influenced by a variety of evolutionary processes. Our simulation-based analyses indicate that *P. tremula* and *P. tremuloides* diverged around 2.2-3.1 Mya during the Late Pliocene and/or Early Pleistocene. This timing corresponds closely with the first opening of the Bering Strait, which occurred 3.1-5.5 Mya and broke up the overland intercontinental migration route of terrestrial floras between Eurasia and North America (Marincovich and Gladenkov 1999; Gladenkov et al. 2002). This may have been less of an immediate barrier to wind-dispersed *Populus* than some other tree species, but the severing of the Bering land bridge associated with the onset of dramatic climatic oscillations throughout the Pleistocene were likely the principal drivers for initial divergence between *P. tremula* and *P. tremuloides* (Comes and Kadereit 1998; Milne and Abbott 2002; Du et al. 2015). Given the modern-day geographic isolation, disjunct distributions and extremely low rates of gene flow, our results support an allopatric model of speciation for these two aspen species (Morjan and Rieseberg 2004). The coalescent-based, intra-specific demographic analyses using MSMC demonstrate that both species have experienced substantial population expansions following long-term declines after species divergence. The population expansion of *P. tremuloides* has occurred over the last 50,000-70,000 years, following the retreat of the penultimate glaciation and has continued up to the present (Kaufman and Manley 2004). *P. tremula*, in contrast, experienced a more extended population contraction and, consistent with many other forest trees in Europe, the initiation of the population expansion in *P. tremula* coincided with the end of the Last Glacial Maximum (Hewitt 2000; Hewitt 2004).

Consistent with the expectation for allopatric speciation, where the absence of gene flow allowed for the accumulation of inter-specific differentiation due to stochastic genetic drift (Coyne and Orr 2004), we detect widespread genomic differentiation between the two species. Although neutral processes are likely responsible for the majority of the genetic differentiation we observe between the two species on a genome-wide scale (Coyne and Orr 2004; Strasburg et al. 2012), a number of regions exhibit more extreme genetic differentiation compared to expectations based on demographic simulations and thses regions also show multiple evidences of the action of natural selection (Nielsen et al. 2009). If natural selection has truly been one of the dominant evolutionary forces shaping genetic differentiation between the two species, regions of low recombination would be expected to show increased *F*_ST_ values, but not increased d_xy_ values (Noor and Bennett 2009; Cruickshank and Hahn 2014). This occurs because natural selection (through either selective sweeps and/or background selection) removes neutral variation over longer distances in regions of low recombination (Begun and Aquadro 1992). As a consequence, relative measures of divergence (e.g. *F*_ST_) that rely on within-species diversity are expected to be higher in regions with restricted recombination (Noor and Bennett 2009; Nachman and Payseur 2012). In contrast, increased absolute divergence (e.g. d_xy_) is only expected if reduced gene flow occurred in regions of low recombination (Nachman and Payseur 2012). In accordance with this view, we observe a significant negative relationship between population-scaled recombination rates (*ρ* and *F*_ST_, but not with d_xy_, in both species (Noor and Bennett 2009; Keinan et al. 2010). Contrary to expectations of heterogeneous gene flow, our findings highlight a significant effect of linked selection in generating the heterogeneous differentiation landscape we observe between the two *Populus* species (Turner and Hahn 2010; Cruickshank and Hahn 2014; Burri et al. 2015).

Rather than being physically clustered into just a few large, discrete genomic ‘islands’ as expected when species diverge in the presence of gene flow (Turner et al. 2005), differentiation islands in our study system are distributed throughout the genome, being narrowly defined and mostly located in regions with substantially suppressed recombination. Combined with the multiple signatures of positive selection we observe in these regions, these islands of divergence likely represent candidate regions harboring loci closely tied to species-specific adaptation rather than loci that are “resistant” to gene flow (Turner and Hahn 2010). In addition, we find no functional over-representation for candidate genes located in these regions, suggesting that a wide range of genes and functional categories have likely been involved in the adaptive evolution of these two species (Wolf et al. 2010).

In addition to the highly differentiated regions that show signs of species-specific positive selection, we also identify a number of lowly differentiated regions that are candidates for being affected by balancing selection in both species (Charlesworth 2006). Apart from low inter-specific divergence and high intra-specific diversity, these regions also contain an excess of sites at intermediate frequencies, a greater proportion of shared polymorphisms between the two species and lack of fixed inter-specific differences. Genes within these regions are enriched for immune and defense response, signal transduction and apoptosis, highlighting the influence of co-evolutionary arms races between hosts and natural enemies on the persistence of functional genetic diversity in immunity and defense-related genes (Tiffin and Moeller 2006; Salvaudon et al. 2008). That said, caution is required when interpreting the functional properties of the regions identified here, and future studies of these candidate genes are clearly needed to better assess the adaptive genetic potential of these widespread forest tree species to current and future climate change.

A number of factors may have influenced our demographic inferences and the detection of natural selection in the two species. First, the presence of within-species population subdivision could have magnified the inference of demographic expansion in *P. tremuloides*, because pooling samples from populations of Alberta and Wisconsin skews the SFS toward low-frequency polymorphism (more negative Tajima’s D) (Figure S4) and results in larger estimates of effective population sizes than estimates obtained from local samples (Figure S3). However, all our analyses suggest that this confounding effect is very weak (see Results) and the divergence between the two subpopulations of *P. tremuloides* likely is too recent to have any major effects on the demographic reconstruction and tests of selection in these species (Chikhi et al. 2010). Another caveat concerns the sampling scheme used in the two species. Local samples in *P. tremula* may not adequately reflect species-wide demography compared to the pooled samples in *P. tremuloides*. However, the extent to which this might influence the estimates of inter-specific *F* ST deserves further study. More generally, sampling should likely be more extensive in both species to capture a greater proportion of the species-wide diversity, although local sampling is expected to only have small effects in species with high gene flow like *Populus* (Wakeley 2000). Finally, inter-specific hybridization in either species could potentially bias our results. However, there are no other species of *Populus* occurring in regions where the *P. tremula* individuals were sampled. For *P. tremuloides*, naturally occurring hybridization is only known to occur with *P. grandidentata* in central and eastern North America where the two species co-occur (Pregitzer and Barnes 1980). Therefore, any possible hybridization in our study would be limited to samples from the Wisconsin population of *P. tremuloides* but, as noted above, we did not detect any major differences in patterns of genetic variation between the two subpopulations, suggesting little or no effect of hybridization.

## Conclusion

Here we provide insights into the speciation and recent evolutionary histories of two closely related forest tree species, *P. tremula* and *P. tremuloides*. Our study supports an allopatric model of speciation for the two species, which are estimated to have diverged around 2.2-3.1 Mya as a result of the first opening of Bering Strait. Coalescent simulations suggest that stochastic genetic drift and historical demographic processes can largely explain the genome-wide patterns of differentiation between species. However, there is an excess of regions displaying extreme inter-specific genetic differentiation in the observed data compared with demographic simulations, which is likely indicative of the action of natural selection. In addition, we find that heterogeneous genomic divergence is strongly driven by linked selection and variation in recombination rate in the two species. Instead of being clustered into a few large genomic “islands” as is expected under a model of speciation with gene flow, regions of pronounced differentiation are characterized by multiple signatures of positive selection in one or both species, and are distributed throughout the genome at many small, independent locations. Regions displaying exceptionally low differentiation are likely candidate targets of long-term balancing selection, which are strongly enriched for genes involved in immune and defense response, signal transduction and apoptosis, suggesting a possible link to long-term co-evolutionary arms races with pest and pathogens. Our study highlights that future work should integrate more information on the natural histories of speciation, such as divergence time, geographical context, magnitudes of gene flow, demographic histories and sources of adaptation, when interpreting the meaning of observed patterns of genomic divergence between closely related species.

## Materials and Methods

## Population samples, sequencing, quality control and mapping

A total of 24 individuals of *P. tremula* and 22 individuals of *P. tremuloides* were collected and sequenced (Figure 1a and Table S1). For each individual, genomic DNA was extracted from leaf samples and paired-end sequencing libraries with an insert size of 600bp were constructed according to the Illumina library preparation protocol. Sequencing was carried out on the Illumina HiSeq 2000 platform at the Science for Life Laboratory in Stockholm, Sweden. All samples were sequenced to a target coverage of 25_X_. The sequencing data has been deposited in the Short Read Archive (SRA) at NCBI under accession numbers SRP065057 and SRP065065 for samples of *P. tremula* and *P. tremuloides*, respectively.

For raw sequencing reads (Wang et al. 2015), we used Trimmomatic (Lohse et al. 2012) to remove adapter sequences and cut off bases from either the start or the end of reads when the base quality was lower than 20. Reads were completely discarded if there were fewer than 36 bases remaining after trimming. We then mapped all reads to the *P. trichocarpa* reference genome (v3.0) (Tuskan et al. 2006) with default parameters implemented in bwa-0.7.10 using the BWA-MEM algorithm (Li unpublished data, http://arxiv.org/abs/1303.3997, last accessed May 26, 2013). Local realignment was performed to correct for the mis-alignment of bases in regions around insertions and/or deletions (indels) using RealignerTargetCreator and IndelRealigner in GATK v3.2.2 (DePristo et al. 2011). In order to account for the occurrence of PCR duplicates introduced during library construction, we used MarkDuplicates in Picard (http://picard.sourceforge.net) to remove reads with identical external coordinates and insert lengths. Only the read with the highest summed base quality was kept for downstream analyses.

## Filtering sites

Prior to variant and genotype calling, we employed several filtering steps to exclude potential errors caused by paralogous or repetitive DNA sequences. First, after investigating the empirical distribution, we removed sites showing extremely low (<100 reads across all samples per species) or high (>1200 reads across all samples per species) read coverage. Second, as a mapping quality score of zero is assigned for reads that could be equally mapped to multiple genomic locations, we removed sites containing more than 20 such reads among all samples in each species. Third, we removed sites that overlapped with known repeat elements as identified by RepeatMasker (Tarailo-Graovac and Chen 2009). After all filtering steps, there were 42.8% of sites across the genome left for downstream analyses. Among them, 54.9% were found within gene boundaries, and the remainder (45.1%) was located in intergenic regions.

## SNP and genotype calling

We employed two complementary approaches for SNP and genotype calling (Figure S1): (i) Direct estimation without calling genotypes was implemented in the software ANGSD v0.602 (Korneliussen et al. 2014). Only reads with a minimal mapping quality of 30 and bases with a minimal quality score of 20 were considered. For all filtered sites in both species, we defined the alleles that were the same as those found in the *P. trichocarpa* reference genome as the ancestral allelic state. We used the - doSaf implementation to calculate the site allele frequency likelihood based on the SAMTools genotype likelihood model at all sites (Li et al. 2009), and then used the - realSFS implementation to obtain a maximum likelihood estimate of the unfolded SFS using the Expectation Maximization (EM) algorithm (Kim et al. 2011). Several population genetic statistics were then calculated based on the global SFS (Figure S1). (ii) Multi-sample SNP and genotype calling was implemented in GATK v3.2.2 with HaplotypeCaller and GenotypeGVCFs (Figure S1) (DePristo et al. 2011). A number of filtering steps were performed to reduce false positives from SNP and genotype calling: (1) Remove SNPs that were located in regions not passing all previous filtering criteria; (2) Removed SNPs with more than 2 alleles in both species; (3) Removed SNPs at or within 5bp from any indels; (4) Assigned genotypes as missing if their quality scores (GQ) were lower than 10, and then removed SNPs with more than two missing genotypes in each species; (5) Removed SNPs showing significant deviation from Hardy-Weinberg Equilibrium (*P*<0.001) in each species.

## Population structure

Population genetic structure was inferred using the program NGSadmix (Skotte et al. 2013), with only sites containing lower than 10% of missing data being used. We used the SAMTools model (Li et al. 2009) in ANGSD to estimate genotype likelihoods and then generated a beagle file for the subset of the genome that was determined as being variable using a likelihood ratio test (*P*-value <10^−6^) (Kim et al. 2011). We predefined the number of genetic clusters *K* from 2 to 5, and the maximum iteration of the EM algorithm was set to 10,000.

As another method to visualize the genetic relationships among individuals, we performed principal component analysis (PCA) using ngsTools accounting for sequencing errors and uncertainty in genotype calls (Fumagalli et al. 2014). The expected covariance matrix across pairs of individuals in both species was computed based on the genotype posterior probabilities across all filtered sites. Eigenvectors and eigenvalues from the covariance matrix were generated with the R function eigen, and significance levels were determined using the Tracy-Widom test as implemented in EIGENSOFT version 4.2 (Patterson et al. 2006).

## Demographic history

We inferred demographic histories associated with speciation for *P. tremula* and *P. tremuloides* using a coalescent simulation-based method implemented in *fastsimcoal* 2.5.1 (Excoffier et al. 2013). Two-dimensional joint site frequency spectrum (2D-SFS) was constructed from posterior probabilities of sample allele frequencies by ngsTools (Fumagalli et al. 2014). 100,000 coalescent simulations were used for the estimation of the expected 2D-SFS and log-likelihood for a set of demographic parameters in each model. Global maximum likelihood estimates for each model were obtained from 50 independent runs, with 10-40 conditional maximization algorithm cycles. Model comparison was based on the maximum value of likelihood over the 50 independent runs using the Akaike information criterion (AIC) and Akaike’s weight of evidence (Excoffier et al. 2013). The model with the maximum Akaike’s weight value was chosen as the optimal one. We assumed a mutation rate of 2.5×10^−9^ per site per year and a generation time of 15 years in *Populus* (Koch et al. 2000) when converting estimates to units of years and individuals. Parameter confidence intervals of the best model were obtained by 100 parametric bootstraps, with 50 independent runs in each bootstrap.

We then employed multiple sequential Markovian coalescent (MSMC) method to estimate variation of scaled population sizes (*N*_e_) over historical time in both species (Schiffels and Durbin 2014), which is an extension of a pairwise sequential Markovian coalescent (PSMC) method (Li and Durbin 2011). Prior to the analysis, all segregating sites within each species were phased and imputed using fastPHASE v1.4.0 (Scheet and Stephens 2006). A generation time of 15 years and a rate of 2.5×10^−9^ mutations per nucleotide per year (Koch et al. 2000) were used to covert the scaled times and population sizes into real times and sizes.

## Genome-wide patterns of differentiation

We have previously shown that linkage disequilibrium (LD) decays within 10 kilobases (Kbp) in both *P. tremula* and *P. tremuloides* (Wang et al. forthcoming), and we thus divided the genome into 39,406 non-overlapping windows of 10 Kbp in size to investigate patterns of genomic differentiation between species. For a window to be included in the downstream analyses, we required there to be at least 1000 bases left after all above filtering steps. Levels of genetic differentiation between species at each site were estimated using method-of-moments *F*_ST_ estimators implemented in ngsFST from the ngsTools package (Fumagalli et al. 2014), which calculates indices of the expected genetic variance between and within species from posterior probabilities of sample allele frequencies, without relying on SNP or genotype calling (Fumagalli et al. 2013). We then averaged *F*_ST_ values across all sites within each 10 Kbp non-overlapping window.

## Coalescent simulations for detecting outlier windows

In order to examine thresholds for detection of outlier windows that may have been targets of natural selection, we conducted coalescent simulations to compare observed patterns of genetic differentiation (*F*_ST_) to those expected under different demographic models (see Results). All simulations were performed using the program *msms* v3.2rc (Ewing and Hermisson 2010) based on demographic parameters derived from the best-fitting model inferred by *fastsimcoa12.5.1* (Excoffier et al. 2013). Population-scaled recombination rates (*ρ*) were assumed to be between 1 Kbp^-1^ and 5 Kbp^-1^ given the large variation we found in both species (Wang et al. forthcoming). We simulated genotypes corresponding to a 10 Kbp region with the same sample size as the real data for 100,000 replications, from where we simulate genotype likelihoods using the program msToGlf in ANGSD (Korneliussen et al. 2014) by assuming a mean sequencing depth of 20X and an error rate of 0.5%. We estimated two summary statistics, nucleotide diversity (θ_π_) and Tajima’s D, from sample allele frequency likelihoods in ANGSD for all simulation replicates to test whether the simulated data matches the observed data. To assess whether any of the observed windows display *F*_ST_ values deviating significantly from neutral expectations, we determined the conditional probability (*P*-value) of observing more extreme inter-specific *F*_ST_ values among simulated data sets than among the observed data. Our significance was based on running 500,000 coalescent simulations of the most acceptable demographic null model (see Results). We then corrected for multiple testing by using the False Discovery Rate (FDR) adjustment, and only windows with FDR lower than 1% were considered as candidate regions affected by selection (Storey 2002).

## Molecular signatures of selection in outlier regions

To assess the occurrence of selection in outlier windows displaying either exceptionally high or low differentiation, we compared these two unions of outlier windows to the remaining portion of the genome by a variety of additional population genetic statistics in both species. First, θ_π_, Tajima’s D (Tajima 1989) and Fay & Wu’s H (Fay and Wu 2000) were calculated from sample allele frequency likelihoods in ANGSD over non-overlapping 10 Kbp windows. Second, levels of LD and population-scaled recombination rates (ρ)were estimated and compared. To evaluate levels of LD within each 10 Kbp window, the correlation coefficients (*r^2^*) between SNPs with pairwise distances larger than 1 Kbp were calculated using VCFtools v0.1.12b (Danecek et al. 2011). Population-scaled recombination rates (*ρ* were estimated using the Interval program of LDhat 2.2 (McVean et al. 2004) with 1,000,000 MCMC iterations sampling every 2,000 iterations and a block penalty parameter of five. The first 100,000 iterations of the MCMC iterations were discarded as burn-in. Resulting estimates of *r*^2^ and ρ were averaged over each 10 Kbp window. In both species, windows were discarded in the estimation of *r*^2^ and *ρ* if there were less than 3 Kbp and/or 10 SNPs left from previous filtering steps. Finally, we used the program ngsStat (Fumagalli et al. 2014) to calculate several additional measures of genetic differentiation: (1) with *P. trichocarpa* as an outgroup, the proportion of fixed differences that is caused by derived alleles fixed in either *P. tremula* or *P. tremuloides* among all segregating sites; (2) the proportion of inter-species shared polymorphisms among all segregating sites; (3) d_xy_, which was calculated from sample allele frequency posterior probabilities at each site and was then averaged over each 10 Kbp window; and (4) the relative node depth (RND), which was calculated by dividing the d_xy_ of the two aspen species with d_xy_ between aspen (represented by 24 samples of *P. tremula* in this study) and *P. trichocarpa* (24 samples; see Wang et al. forthcoming). Significance of the differences between outlier windows and the genome-wide averages for all above mentioned population genetic statistics were examined using one-sided Wilcoxon ranked-sum tests.

## Gene ontology (GO) enrichment

To determine whether any functional classes of genes were overrepresented among regions that were candidates for being under selection, we performed functional enrichment analysis of GO using Fisher’s exact test by agriGO’s Term Enrichment tool (http://bioinfo.cau.edu.cn/agriGO/index.php) (Du et al. 2010). GO groups with fewer than two outlier genes were excluded from this analysis. *P*-values of Fisher’s exact test were further corrected for multiple testing with Benjamini-Hochberg false discovery rate (Benjamini and Hochberg 1995). GO terms with a corrected *P*-value <0.05 were considered to be significantly enriched.

## Acknowledgements

We are grateful to Rick Lindroth for providing access to the samples of *P. tremuloides* used in this study. We thank Carin Olofsson for extracting DNA for all samples used in this study. We thank both the editor and two anonymous referees for their useful comments on the manuscript. The research has been funded through grants from Vetenskapsrådet and a Young Researcher Award from Umeå University to PKI. JW was supported by a scholarship from the Chinese Scholarship Council. The authors also would like to acknowledge support from Science for Life Laboratory, the National Genomics Infrastructure (NGI), and Uppmax for providing assistance in massive parallel sequencing and computational infrastructure.

